# Coadaptation of coexisting plants enhances productivity in an agricultural system

**DOI:** 10.1101/2023.02.08.527628

**Authors:** Anja Schmutz, Christian Schöb

## Abstract

- Growing crops in more diverse crop systems (i.e. intercropping) is one way to produce food more sustainably. Even though intercropping, compared to average monocultures, is generally more productive, the full yield potential of intercropping might not yet have been achieved as modern crop cultivars are bred to be grown in monoculture. Breeding plants for more familiarity in mixtures, i.e. plants that are adapted to more diverse communities (i.e. *adaptation*) or even to coexist with each other (i.e. *coadaptation*) might have the potential to sustainably enhance productivity.
- In this study, the productivity benefits of familiarity through *evolutionary adaptation*, where one species adapts to its neighbourhood, and *coevolutionary coadaptation*, where two or more species adapt to each other, were disentangled in a crop system through an extensive common garden experiment. Furthermore, evolutionary and coevolutionary effects on species-level and community-level productivity were linked to corresponding changes in functional traits.
- We found evidence for higher productivity and trait convergence with increasing familiarity of the plants composing the community. Furthermore, our results provide evidence for *coevolution* of plants in mixtures leading to higher productivity of coadapted species. However, with the functional traits measured in our study we could not fully explain the productivity benefits found upon *coevolution*.
- Our study is, to our knowledge, the first study that investigated *coevolution* among randomly interacting plants and was able to demonstrate that *coadaptation* through *coevolution* of coexisting species in mixtures promote ecosystem functioning (i.e. higher productivity). This result is particularly relevant for the diversification of agricultural and forest ecosystems, demonstrating the added value of artificially selecting plants for the communities they are familiar with.

## Introduction

Agriculture is under pressure to produce food more sustainably (FAO, 2018). Growing crops in mixtures (called intercropping or mixed cropping) is an often proposed method for a sustainable intensification of agriculture (Lithourgidis et al., 2011; Martin-Guay et al., 2018). Mixed cropping can maintain similar yields to monocropping on less land area and with less fertiliser inputs (Li et al., 2023, 2020). These yield benefits of mixtures compared to monocultures can be attributed to a range of mechanisms, such as niche complementarity in resource use (Engbersen et al., 2021; Schmutz and Schöb, 2022) or stimulation of plant growth-promoting microbes (Stefan et al., 2021) that exploit complementarity of the different crop species in mixtures (Brooker et al., 2015; Engbersen et al., 2022).

Even though mixed cropping generally produces more yield per unit land area than monocropping, the full yield potential of mixed cropping might not yet have been reached. Most of the commonly grown crops nowadays are bred for and multiplied in monocultures where they experience only intraspecific interactions and are thus not necessarily well suited to be grown in mixtures where also interspecific interactions occur. Hence, crop ecotypes specifically bred for mixtures might achieve additional yield benefits (Bourke et al., 2021; Brooker et al., 2021; Chen et al., 2021; Moore et al., 2022). Indeed, ecological studies conducted in grasslands have shown that plants adapt to growth in polycultures, with significant positive effects on community-level productivity (Zuppinger-Dingley et al., 2014) due to less intense competition in mixtures of adapted plants compared to the same mixtures of naïve plants (Schöb et al., 2018) – a result that could be confirmed in an annual crop system (Stefan et al., 2022).

Plants that share a common coexistence history can familiarise themselves and adapt to each other within a very short time frame (Thorpe et al., 2011). However, whether this adaptation is the result of *coevolution* where the interacting agents co-adapt in a specific manner to increase complementarity or whether this adaptation is the result of *evolution* of a general mixture ecotype that performs better with heterospecific neighbours than a monoculture ecotype, is to our knowledge not known. Coevolution of plants with other clades such as insects (i.e. pollinators or parasites) or animals (i.e. pollinators or dispersers) are well studied (Cogni et al., 2022). In contrast, the coevolution among plants, at least when there is no obligate interaction such as an obligate parasitism, received much less attention (Thorpe et al., 2011). In general, coevolution is suggested to play a key role in diversification and speciation (Thompson, 2016, 2012) and is important for ecosystem functioning (Loeuille et al., 2002). Therefore, if plants coevolved, we would expect them to cause a positive effect on ecosystem functioning (i.e. higher productivity).

Enhanced productivity in plant communities is mainly driven by differences in resource use strategies (Callaway et al., 2003). Therefore, shifts to different resource use strategies of coexisting species (i.e. character displacement) upon their interaction can explain enhanced productivity in mixtures (Beans, 2014). If this interaction is very persistent, this character displacement can become adaptive and be inherited to forthcoming generations, leading the traits of the coexisting species to diverge (Grime, 2006). Indeed, trait divergence among coevolved grassland species has been attributed to the corresponding productivity benefits (Zuppinger-Dingley et al., 2014).

To test whether an observed adaptation of coexisting plants is the result of *coevolution* or *evolution*, is an interesting question per se, but particularly relevant when it comes to the selection of community (i.e. monoculture or mixture) ecotypes of crops or breeding programs for community-type cultivars. If there is a general *adaptation* to growth in community (i.e. *evolution*), the selection of a generic community-type cultivar will suffice, while specific *coadaptation* of crop species or even cultivars (i.e. *coevolution*) would require specific cultivars for each of the different communities that would like to be improved.

Here, we used seeds from six crop species that were either grown in community with the specific species for three years or seeds from single plants that had no coexistence history and were therefore naïve to any interaction with other plants. By growing plants with or without common coexistence history in all possible combinations, we were able to test if plants that had a common coexistence adapted to their neighbours. Particularly, the following research questions were addressed: Does adaptation of coexisting plants increase community productivity? Do plants coevolve and show specific *coadaptation* to interacting plants or do plants evolve a generic *adaptation* to the community? What are the consequences of *coadaptation* and *adaptation* on ecosystem functioning? Are *coadaptation* and *adaptation* subject to trait convergence or divergence?

These questions were addressed on the community level (**hypotheses 1** and **6**) and the species level (**hypotheses 2-5**) (**Fig. 1**). We hypothesised that community productivity increases with increasing level of familiarity of coexisting species (**hypothesis 1, Fig. 1a**). Furthermore, we assumed that plants with coexistence history grown together are more productive than plants with no coexistence history (*adaptation* hypothesis, **hypothesis 2, Fig. 1b**). If this adaptation is a result of *coevolution*, we would expect that plants with coexistence history grown with “known” plants produce more than plants with coexistence history that grew with “unknown” plants (*coevolution* hypothesis, **hypothesis 3, Fig. 1b**). Furthermore, if adaptation among coexisting plants is a result of *evolution*, we assumed that in a community with both coexisting and naïve plants, the coexisting plant would produce more while the naïve plant of the same community would not (*evolution* hypothesis, **hypothesis 4, Fig. 1b**). We would expect this as coexisting plants are “accustomed” to interactions in a community with homo- and heterospecific neighbours whereas naïve plants are not.

**Fig. 1.**
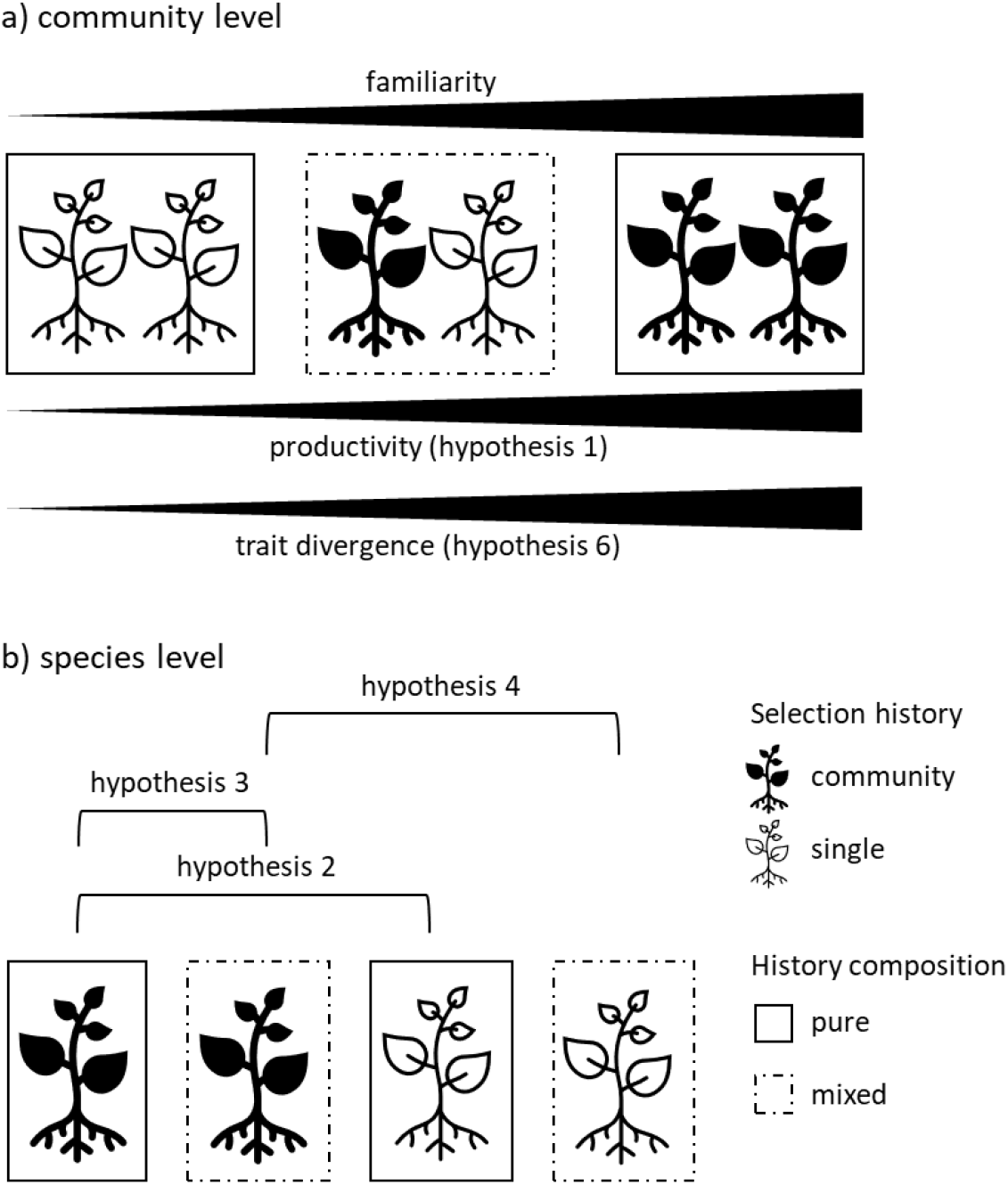
Conceptual framework of the study at (**a**) community and (**b**) species level, with the pairwise contrasts to test specific hypotheses. The *community* selection history are plants with a coexistence history (coexisting plants), *single* selection history are plants with no coexistence history (naïve plants). These plants were either grown in *pure* (only *community* or *single* selection history) or in *mixed* history composition (*community* and *single* selection history grown together).

Besides plant productivity, different functional traits such as plant height, specific leaf area (SLA) and leaf dry matter content (LDMC) were measured. These functional traits capture different resource use strategies where plant height is associated with light use, SLA with nutrient use and growth strategy and LDMC with water use and leaf toughness (Pérez-Harguindeguy et al., 2013). Hence, we would expect that changes in productivity as a result of adaptation (hypotheses 2-4) are due to changes in functional traits – and therefore changes in resource use strategy (**hypothesis 5, Fig. 1b**). Additionally, we assumed that increasing productivity with increasing familiarity (**hypothesis 1**) is a result of increasing trait divergence (**hypothesis 6, Fig. 1a**).

## Material & Methods

### Experimental site

The experiment was conducted in an outdoor common garden at the research station Aprisco in Torrejón el Rubio, Cáceres, Spain (39°48’48”N 6°00’01”W). The region has a Mediterranean climate with dry, hot summers and wet, mild winters. The weather during the duration of the experiment (February until July 2021) was characterised by mild to hot temperatures and a few rain events (**Fig. 2**). The experimental garden included 1600 square plots that measured 25×25 cm. The plots were further arranged in beds of 1×10 m composed of 4×40 plots. Cultures were randomly allocated within the beds. Throughout the whole growing season, the field was irrigated with an automated irrigation system which was set to keep soil moisture between 50% and 75% of field capacity. During the whole season, plots were weeded regularly to minimise interactions with weeds. Throughout the experiment, no fertiliser was applied to the plots. The soil consisted of approx. 50%, sand 50% and <2% clay, had a pH of 5, and nitrogen and carbon content were <0.1% and 1%, respectively.

**Fig. 2.**
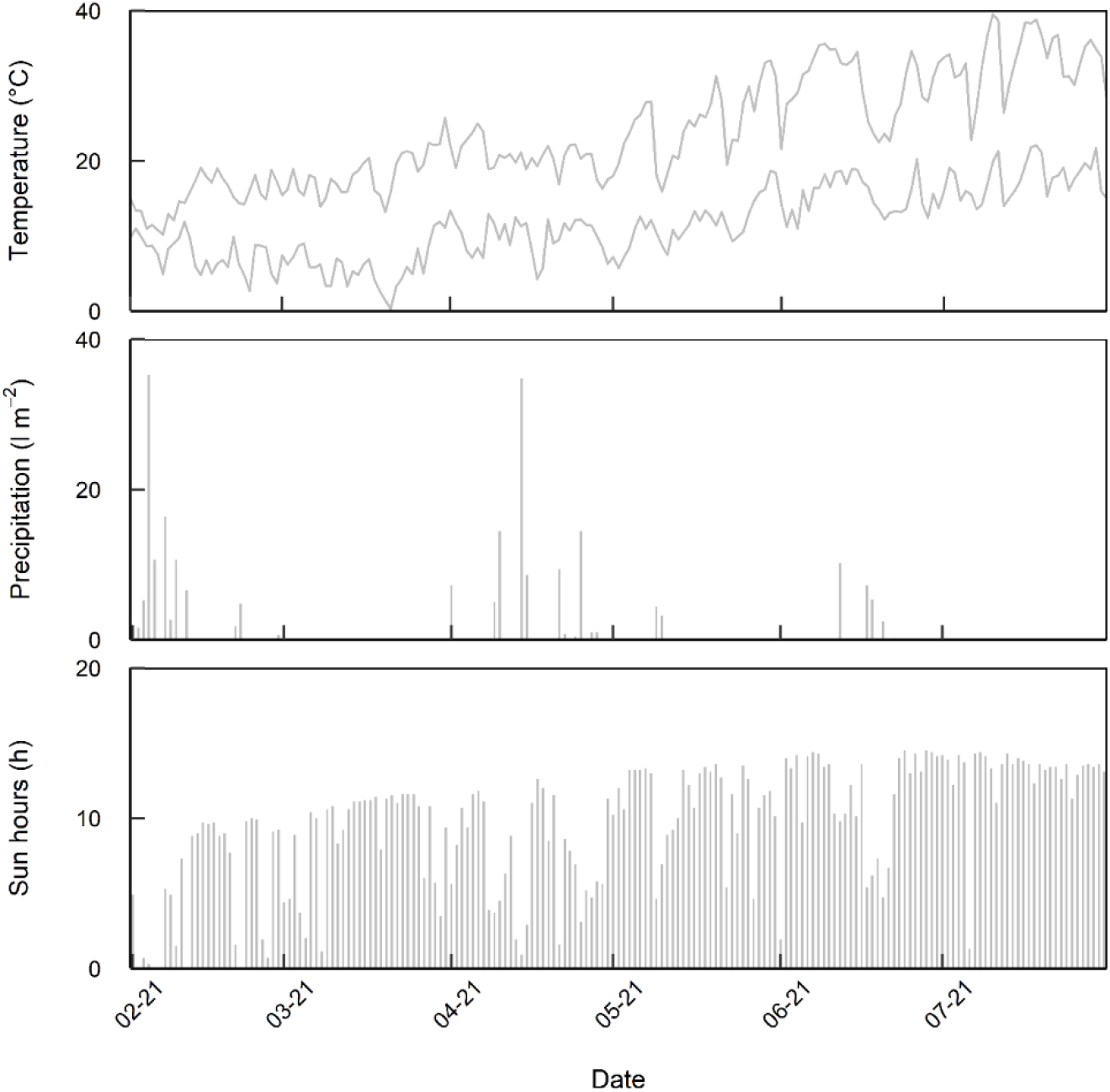
Weather data for the duration of the experiment. (A) Daily maximum and minimum temperature. (B) Daily cumulative precipitation. (C) Daily cumulative sun hours. Data is from a weather station 50 km away. Source: datosclima.es

### Experimental design

For this experiment, six species were used including the cereals oat (*Avena sativa* var. Previsión) and summer wheat (*Triticum aestivum* var. Cabezorro), the legumes lentil (*Lens culinaris* var. de la Armuña) and lupin (*Lupinus angustifolius* wild type) and the herbs camelina (*Camelina sativa* n.a.) and coriander (*Coriandrum sativum* wild type). All six crop species had either no coexistence history (seeds from single plants; *single* selection history) or a common coexistence history (seeds from 2-species and 4-species mixtures; *community* selection history) for three years. In the previous years, seeds from the specific cultures were collected and resown in the same culture the following year (Stefan et al., 2022).

All plants were grown in 2×2 Latin squares (**Table S1**). Four individuals from either the same species (i.e. monoculture) or with a species from a different functional group (i.e. mixture) were grown together (**Fig. S1**). These plots were either composed only with *single* selection history or with *community* selection history (*pure* history composition), or both *single* and *community* selection history (*mixed* history composition) in all possible combinations. In total, the plots with specific species composition and history composition were replicated six times.

Sowing was conducted by hand between 5 and 9 February 2021. For this, one seed per position in the 2×2 Latin square was sown (**Fig. S1**). Additional seeds were sown in empty plots and trays for transplantations in case of death or no germination. Due to poor germination of camelina and attacks by larvae of lupin, these species were resown starting 3 March 2021. For the other species, lacking plants were transplanted between 24 March and 2 April 2021. In some cases (especially lentil and coriander), the number of replicates was reduced to complete plots. Transplantation (yes or no) and age of plants (first or second sowing) was reported for each plant individual.

### Sample collection

Functional traits sampling was undertaken when the first plants were flowering (between 3 and 11 May 2021). We sampled always one individual per position in each plot (**Fig. S1**). Plant height (cm) was measured from the ground up to the base of the last leaf. For specific leaf area (SLA) and leaf dry matter content (LDMC), one leaf per plant was collected and stored in a plastic bag together with wet cotton overnight. The next day, saturated leaves were weighed (fresh leaf weight), scanned (leaf area) and subsequently dried and weighed again (dry leaf weigh). SLA (m^2^ kg^-1^) was calculated by dividing the leaf area (m^2^) by the dry leaf weight (kg); LDMC (mg g^- 1^) by dividing leaf dry weight (mg) by leaf fresh weight (g).

Plants were harvested when fruits were ripe (starting 7 June 2021). Each plant in the plot was harvested separately and separated into aboveground biomass and seeds. Aboveground biomass was dried for 72h at 80°C, seeds were air-dried. Subsequently, (aboveground) biomass and seed mass per plant were weighed and reported. Hence, data was available either at individual level (biomass and seed mass) or at position level (plant height, SLA and LDMC) (**Fig. S1**).

### Analysis

At the very beginning, biomass, seed mass, plant height, SLA and LDMC were checked for normal distribution and subsequently log-transformed (biomass, seed mass, SLA and LDMC) or square-root transformed (plant height). Outliers were identified and removed based on the z-score (outlier if z-score>3). Outliers were only present in the variables that were calculated from measured variables (45 values for SLA and 55 for LDMC).

To investigate evolutionary processes in a plant community, seed mass (i.e. fecundity) is often considered a measure of fitness (i.e. number of offspring in the next generation) (Primack and Kang, 1989). The seed mass data in this experiment were strongly zero-inflated due to low productivity of some individuals, making analyses challenging. Hence, a different approach was used for seed mass (see below). Nevertheless, for annual crop plants it is suggested that biomass and fecundity are related and, thus, biomass might be a reliable proxy for fitness (Younginger et al., 2017). Seed mass and biomass were indeed positively correlated in this experiment (**Fig. S2**), thereby justifying the use of biomass as a response variable in our study. Nevertheless, we did not want to exclude seed mass from the analysis and hence applied a generalised linear mixed model (**Fig. S3b**) with binomial distribution, with the binominal seed mass as response variable (0=no seeds, 1=seeds). The explanatory variables were transplanted (yes vs no), sowing (first vs second), culture (monoculture vs mixture), species (oat, wheat, lentil, lupin, camelina or coriander), selection history (single vs community) and history composition (pure vs mixed), and all possible interactions between species, selection history and history composition. The random terms were the position within plot, the species composition, and the interaction species composition × selection history. The results indicated no effect of adaptation on the outcome of having or not having seed mass (**Table S2**). Hence, we concluded that it is feasible to continue analysis of seed mass to test for adaptation with zero-truncated data.

Harvest index (HI) was calculated from seed mass (zero-truncated) and biomass (HI = seed mass / (biomass + seed mass)). For the functional traits (plant height, SLA and LDMC), a principal component analysis (PCA) was conducted. The first PCA explained 67% of variation and had an eigenvalue > 2 (**Fig. S4**) and thus was used in the subsequent models.

To test for community adaptation (**hypothesis 1**), the plot biomass and plot seed mass (zero-truncated) (sum of the positions a and b) was calculated from the mean of each of the two positions. Furthermore, factor levels that combine the status of transplantation and sowing, respectively, of the two individuals for each position were implemented (e.g. a factor level of “1” indicated that both individuals were not transplanted, a factor level of “1.5” indicates that one individuals was transplanted and the other not, and a factor level of “2” indicates that both individuals were transplanted). Similarly, to test for trait divergence (**hypothesis 6**), the absolute difference in functional trait values (scaled by species) was calculated for each plot and each functional trait. Only cases were both positions had a value were used for further analysis.

Afterwards, linear mixed models (**Fig. S3a**) were conducted with the response variables plot biomass (square-root transformed), seed mass (log-transformed, zero-truncated) and difference in functional traits (square-root transformed, for all functional traits), respectively. The explanatory variables were transplanted (for each position), sowing (for each position), culture (i.e. monoculture vs mixture), and familiarity (numeric; pure *single*=1, *mixed*=2, pure *community*=3). In seed mass, the second-degree polynomial of familiarity was included also as explanatory variable. The random term was the species composition. The level of familiarity starts with a combination of unfamiliar components (pure *single*, i.e. both positions occupied by plants from *single* selection history), over a combination of an unfamiliar with a familiar component (*mixed*, i.e. one position occupied by a plant from *single* selection history and the other position occupied by a from *community* selection history) to a combination of familiar components (pure *community*, i.e. both positions occupied by plants from *community* selection history).

To test the hypotheses at species level (**hypotheses 2-5**), further LMMs were computed. The response variable was either biomass (log-transformed), seed mass (log-transformed, zero-truncated), harvest index (square-root transformed) or coordinates of PC1 from the functional traits (no transformation). The explanatory variables were transplanted (yes vs no), sowing (first vs second), culture (monoculture vs mixture), species (oat, wheat, lentil, lupin, camelina or coriander), selection history (*single* vs *community*) and history composition (*pure* vs *mixed*), and all possible interactions between species, selection history and history composition. The random terms were the plot, the position within plot, the species composition, and the interaction species composition × selection history. For the data that was only available at the position level (PC1 of functional traits), the random term position within plot was dropped, with plot remaining as random term (**Fig. S3b**). Subsequently, analysis of variances (ANOVA) and post-hoc analyses (*estimated marginal means* aka *least-squares means*) were performed to test the hypotheses. Contrast analyses were always conducted between *community* and *single* selection history in both *pure* history composition (**hypothesis 2**), *community* selection history from *pure* history composition and *community* selection history from *mixed* history composition (**hypothesis 3**), and *community* and *single* selection in *mixed* history composition (**hypothesis 4**). Contrasts were compared from back-transformed estimated marginal means.

## Results

### Adaptation and trait convergence/divergence with increasing familiarity (community level)

To test for adaptation and trait divergence at community level (**Fig. 1a**), the relationship of familiarity with biomass and seed mass (**hypothesis 1**) and the functional traits plant height, SLA and LDMC (hypothesis 6) were estimated. With increasing familiarity, the community biomass increased (**Fig. 3a, Table S3**). Communities that consisted purely of individuals which have a common coexistence history had on average approx. 34% more biomass compared to communities with no coexistence history (pure *single* selection history), while communities with one familiar and one unfamiliar component had approx. 20% more than the pure *single* selection history. Seed mass showed similar trends as biomass (**Fig. S5**) – but the second-degree polynomial of familiarity was only marginally significant (**Table S3**).

**Fig. 3.**
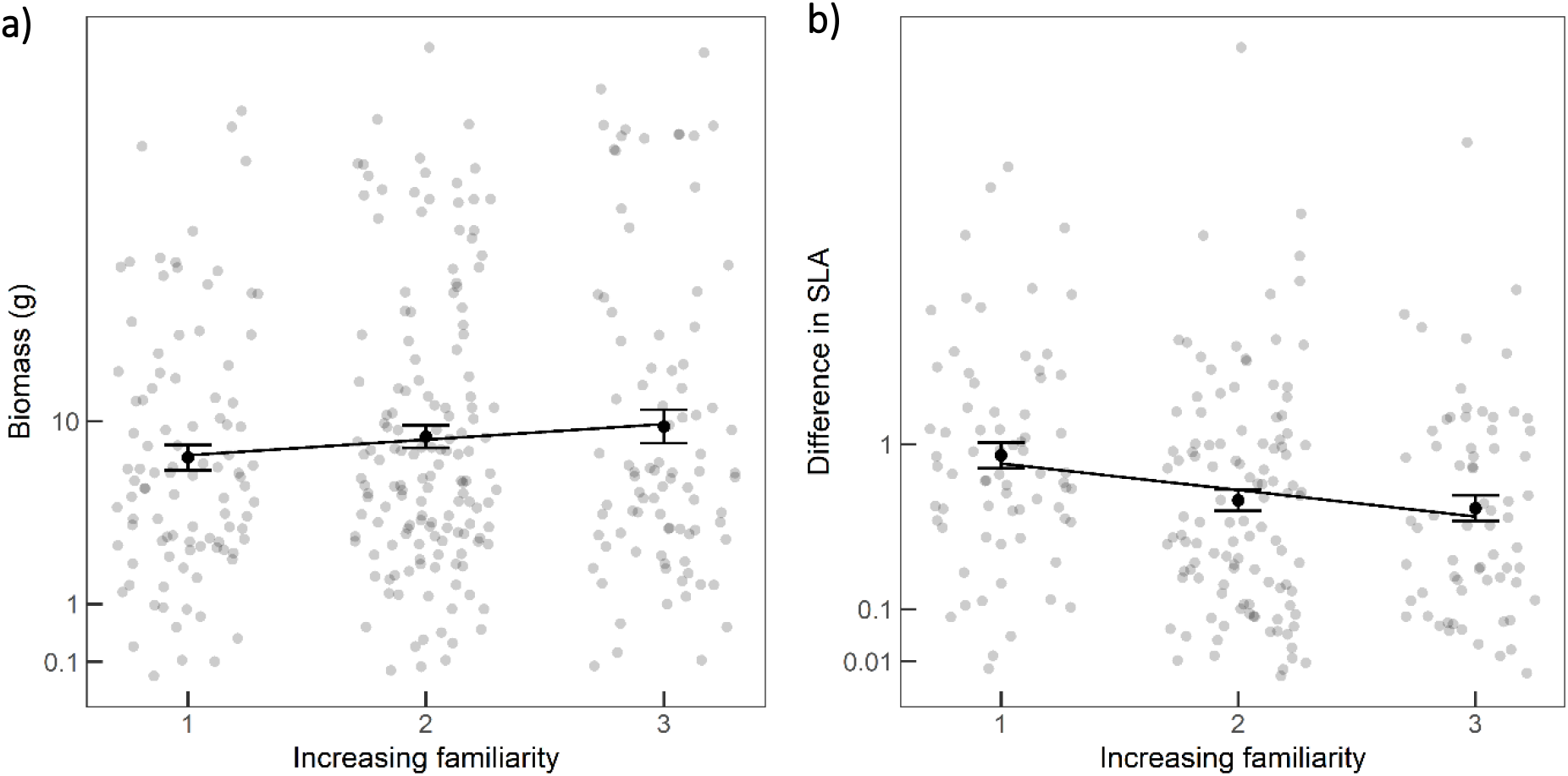
(**a**) Plot biomass (square-root scale) and (b) difference in SLA (square-root scale) of communities with increasing familiarity (pure *single*=1, *mixed*=2, pure *community*=3). Shown are the single data points, the mean ± standard error of the mean and the regression line (formula: y∼x). For statistics see Table S3 and S4.

From the three functional traits, only SLA showed a response to increasing familiarity (**Fig. 3b, Table S4**). Generally, differences in SLA were highest in communities that consisted purely of *single* selection history and converged over *mixed* communities to communities with purely *community* selection history.

### Adaptation and character displacement at species level

To test for adaptation and character displacement at species level (Fig. 1b), contrast analyses for biomass and seed mass (hypotheses 2-4) and for the PC1 of the functional traits (hypothesis 5) were conducted.

With regard to biomass and seed mass, only coriander showed a response to adaptation. Coriander biomass of communities with a pure *community* selection history was significantly (1.5 times) higher than biomass from coriander grown in communities that consisted only of *single* selection history. Furthermore, coriander *community* selection history had a significantly (3.7 times) higher biomass when grown with coexisting plants (*pure* history composition) than when grown in *mixed* history composition (**Fig. 4a**). Seed mass (zero-truncated) partly supported this finding but with weaker differences (only 1.7 times higher seed mass in coriander *community* selection history when comparing *pure* with *mixed* history composition) (**Fig. 4b**). Whether the plants generally had or did not have seeds (binominal seed mass model) was not affected by adaptation (**Table S2**). For the HI, the only significant difference was in wheat between *community* and *single* selection history from both *pure* history composition (1.4 times higher in *community* selection history) (**Fig. S6**). These results suggest that adaptation of coexisting plants is very species-specific (**Table S5**).

**Fig. 4.**
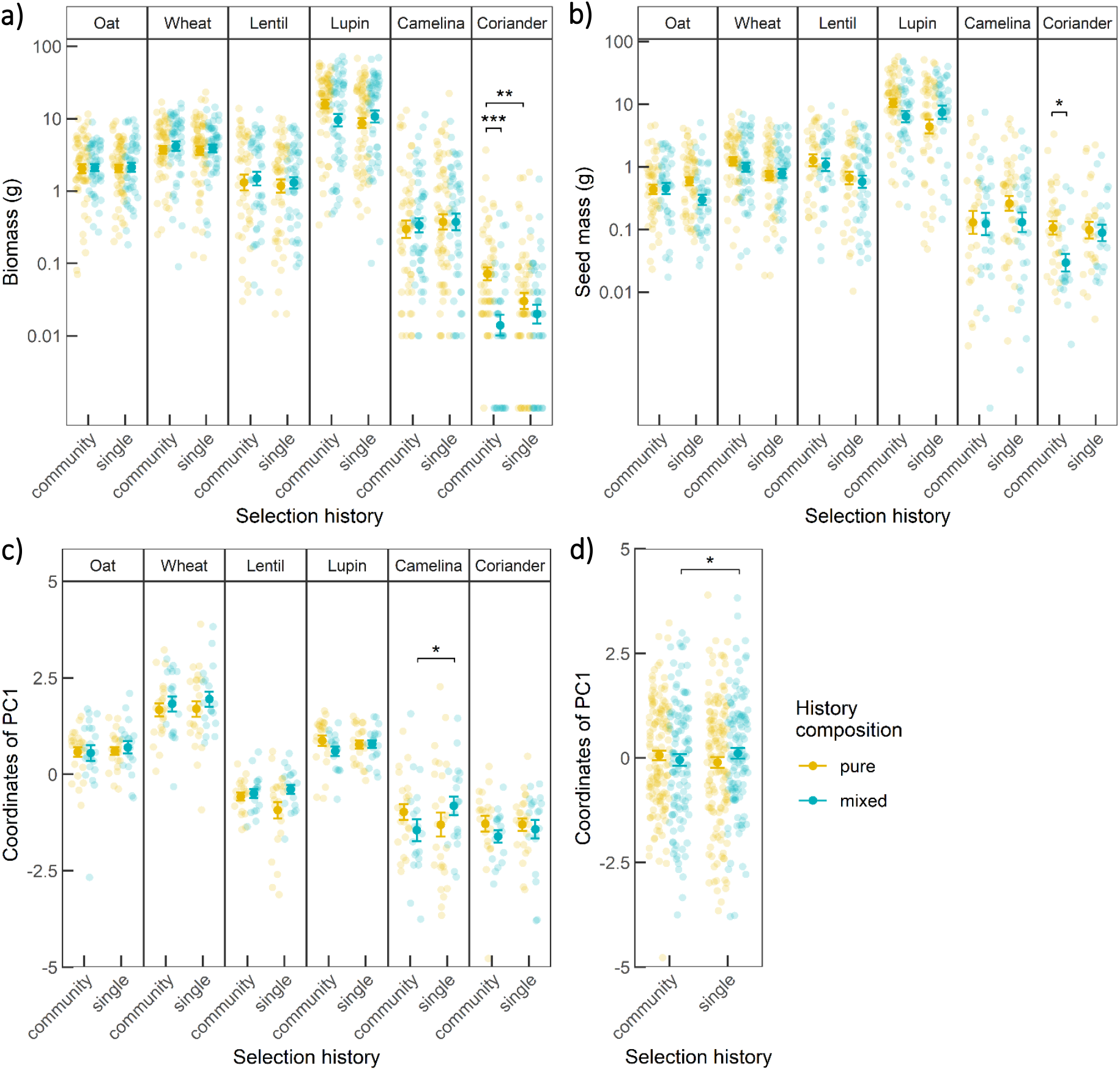
(**a**) Biomass (log-scale), (**b**) seed mass (log-scale, zero-truncated) and PC1 of the functional traits at (**c**) species level and (**d**) across all species of plants from either *community* or *single* selection history and planted in either *pure* (yellow) or *mixed* (blue) history composition. Shown are the single data points and the mean ± standard error of the mean (point and error bar). Significant differences to test the hypotheses (Fig. 1b) are indicated with brackets and asterisk (post-hoc, estimated marginal means). Asterisks correspond to *: P<0.05; **: P<0.01; ***: P<0.001. For statistics see Table S5 and S6.

Concerning the functional traits, a significant difference was present between co*mmunity* and *single* selection history when grown together (*mixed* history composition) at both species level for camelina (**Fig. 4c**) and across all species (**Fig. 4d**). Plants from *community* selection history were generally smaller, had lower SLA and higher LDMC than *single* selection history (**Table S6, Fig. S4**).

## Discussion

We found evidence of increased community productivity with increasing level of familiarity (**hypothesis 1**). Productivity increased from communities that only consisted of plants with no coexistence history, over mixed communities to communities only consisting of plants with a common coexistence history. Contrary to what we hypothesised, we observed trait convergence with increasing familiarity (**hypothesis 6**). This implies that there is a discrimination in selection of plants which would lead to evolutive exclusion of some species.

At the species level, we were able to test if plants adapt to coexisting plants and if this adaption is a result of *evolution* and/or *coevolution*. As far as we know, no other study ever tried to partition these two mechanisms. Results of adaptation were only present in coriander biomass (and partly seed mass). These results suggest *coevolution* rather than *evolution* of coexisting coriander (**hypothesis 3**). Functional traits did not provide evidence (i.e. character displacement) in response to *coevolution* (**hypothesis 5**) but significant changes upon *evolution*.

### Coadaptation of coexisting plants enhances ecosystem functioning

At the community level, we found evidence for increased productivity with increasing familiarity of plant communities. Hence, communities that share (or partly share) a common coexistence history were generally more productive than communities that did not share a common coexistence history. At the species level, we were able to test for adaption among coexisting plants and to distinguish if this adaptation is the result of *evolution* and/or *coevolution*. As hypothesised, the comparison between pure coexisting and pure naïve plant communities can only test for adaptation, which includes both *evolution* and *coevolution*. To examine if this adaptation is the result of *evolution* and/or *coevolution*, other combinations need to be tested. In our study, from a total of six crop species only coriander showed a response upon adaption. Results further suggest that adaptation of coriander to coexisting plants was a result of *coevolution* rather than *evolution*. This finding was supported by both biomass and seed mass.

The harvest index gives indication how resource are allocated in the aboveground plant parts into vegetative and reproductive biomass (Chen et al., 2021; Donald, 1968). Changes towards a higher harvest index would be relevant for the seed crops tested here as plants allocate more resources towards reproductive biomass relative to the vegetative biomass. In this study, we indeed found evidence for a higher harvest index in wheat plants that were grown in communities which share a common coexistence history. This suggests that the hypothesised negative effect of modern crop cultivars on the harvest index in mixtures (Chen et al., 2021) could indeed be reversed with cultivars adapted to grow in mixtures.

In general (at community and at species level), results always indicated a higher productivity due to *coadaptation*. Hence, *coadaptation* of crop species contributes positively to ecosystem functioning (i.e. productivity). This observation is in line with previous research showing that a common coexistence history leads to *adaptation* in mixtures and both higher productivity (Stefan et al., 2022; van Moorsel et al., 2018b, 2018a) and yield stability (van Moorsel et al., 2020). Nevertheless, these studies did not differentiate between *adaptation* and *coadaptation*. Hence, here we are the first to disentangle the effect of *evolution* and *coevolution* of coexisting plants and that *coevolution* between randomly interacting plants occurs and contributes to ecosystem functioning.

Indications for *(co)adaptation* in the investigated crop species was rather low (one species out of six). One reason could be that domestication might have reduced phenotypic plasticity of crops (Semchenko and Zobel, 2005; Vilela and González-Paleo, 2015). But for adaptation to occur, plasticity is essential (Sultan, 1995). Phenotypic plasticity is genetically encoded (Laitinen and Nikoloski, 2019) and, hence, can also be selected for or against (e.g. during breeding/domestication) (Brooker et al., 2022). Breeding has usually focused on plant traits that were relevant for higher yield (Nicotra et al., 2010) and other traits (e.g. relevant for interactions) were generally neglected or even unintentionally selected against (Macfadyen and Bohan, 2010). Furthermore, domestication had large effects on plant physiological mechanisms (Milla et al., 2015) which also changed how crops use resources (Matesanz et al., 2010; Milla et al., 2014). In the last years, breeding for more diverse crop systems (i.e. mixtures) has gained more attention (Bourke et al., 2021; Haug et al., 2021; Moore et al., 2022) and genes were identified which might be relevant for breeding in more diverse crop systems (Wuest et al., 2022; Wuest and Niklaus, 2018). Furthermore, a recent study found evidence for genetic and epigenetic evolution of monoculture and mixture ecotypes in grassland species (van Moorsel et al., 2019). This indicates that a persistent interaction among plants over a certain time has the potential to also alter genes.

Another reason for the limited effects of *(co)adaptation* in our study might be that the three years of coexistence history might not have been long enough for some species to (co)adapt – and especially that this adaptation results in higher yield (Stefan et al., 2022). It is probably more likely that certain traits were favoured that did not lead to higher productivity but were still important for adaptation (Thompson, 2009).

### Character displacement of functional traits upon evolution

We only found evidence for character displacement upon *evolution* in camelina and across all species. This indicates that coexisting and naïve plants grown together had different functional traits, and thus use resources differently. Even though we found evidence for *coevolution* in coriander productivity, we did not find such an indication for *coevolution* in functional traits. So why did the trends we observed in productivity indices not match the trends in the functional traits? Functional traits were correlated with biomass in this study (Fig. S7) and some traits (plant height and SLA) are suggested to show plasticity in response to neighbouring plants (Abakumova et al., 2016). This indicates that the measured traits are indeed relevant for productivity but at the same time are also plastic in response to plant identity. Hence, it might be that *coevolution* is not acting on these specific functional traits (Thompson, 2009) and/or the time was too short for *coevolution* to act on these traits (as also discussed above). On the other hand, the observed character displacement upon *evolution* in SLA is also supported by another study in grassland species (Zuppinger-Dingley et al., 2014). The same study also found character displacement in plant height, which we did not find here. In general, detecting character displacement in plants is more challenging than in other clades since changes in functional traits due to *evolution* are more subtle (Beans, 2014).

Since the study of coevolution among plants is still in a very early stage, it would be interesting to investigate different functional traits to understand on which functional traits *coevolution* is acting. Along this line, plasticity of functional traits in crops is inevitable (Beans, 2014; Sultan, 1995). To set the focus also on root functional trait and its plasticity might be a good start (Schneider and Lynch, 2020). We could previously show that plants show plasticity of water uptake in response to plant diversity (Schmutz and Schöb, 2022) – and root traits have the capability to evolve in response to nutrient availably (Grossman and Rice, 2012).

### Trait convergence and competitive exclusion

Contrary to what we hypothesised we found trait convergence rather than trait divergence of SLA in communities with increasing familiarity at the community level. As suggested by theory, plants that share a limiting resource usually show trait convergence, whereas in a system, where plants have distinct limiting resource, traits are more probable to converge (Slatkin, 1980). If plants use the same resources and their traits are converging, competition between species is intensified and might even lead to competitive exclusion (Gause, 1934; Hardin, 1960). In comparison, trait divergence reduces competition and might result in a stable coexistence (Lawlor and Smith, 1976). These mechanisms, which are already discussed in more detail elsewhere (McPeek, 2019; Pásztor et al., 2020), can also be applied to our (agricultural) system. The fact that with increasing familiarity productivity increased and SLA converged (**Fig. 3**), and that productivity and SLA were negatively related (**Fig. S7**) implies that more productive plants with similar (and thus low) SLA are “favoured” upon *coadaptation*. Hence, there is a discrimination in selection of plants in our system which leads to the evolutive (i.e. over multiple generations) exclusion of plants with lower productivity and higher SLA (**Fig. 5**, scenario A). In the alternative (hypothetical) scenario with trait divergence instead of convergence, more productive plants with distinct (which include low and medium) SLA are “favoured” (**Fig. 5**, scenario B). This would still lead to the evolutive exclusion of plants with very low productivity – but since distinct SLA are also favoured, plants with high (or medium) productivity and distinct SLA would be selected for. This implies that, when breeding/selecting for coadapted plant ecotypes, it is necessary to also consider differences in functional traits (i.e. trait divergence) to ensure that coadapted ecotypes have an evolutive (over multiple generations) stable productivity.

**Fig. 5.**
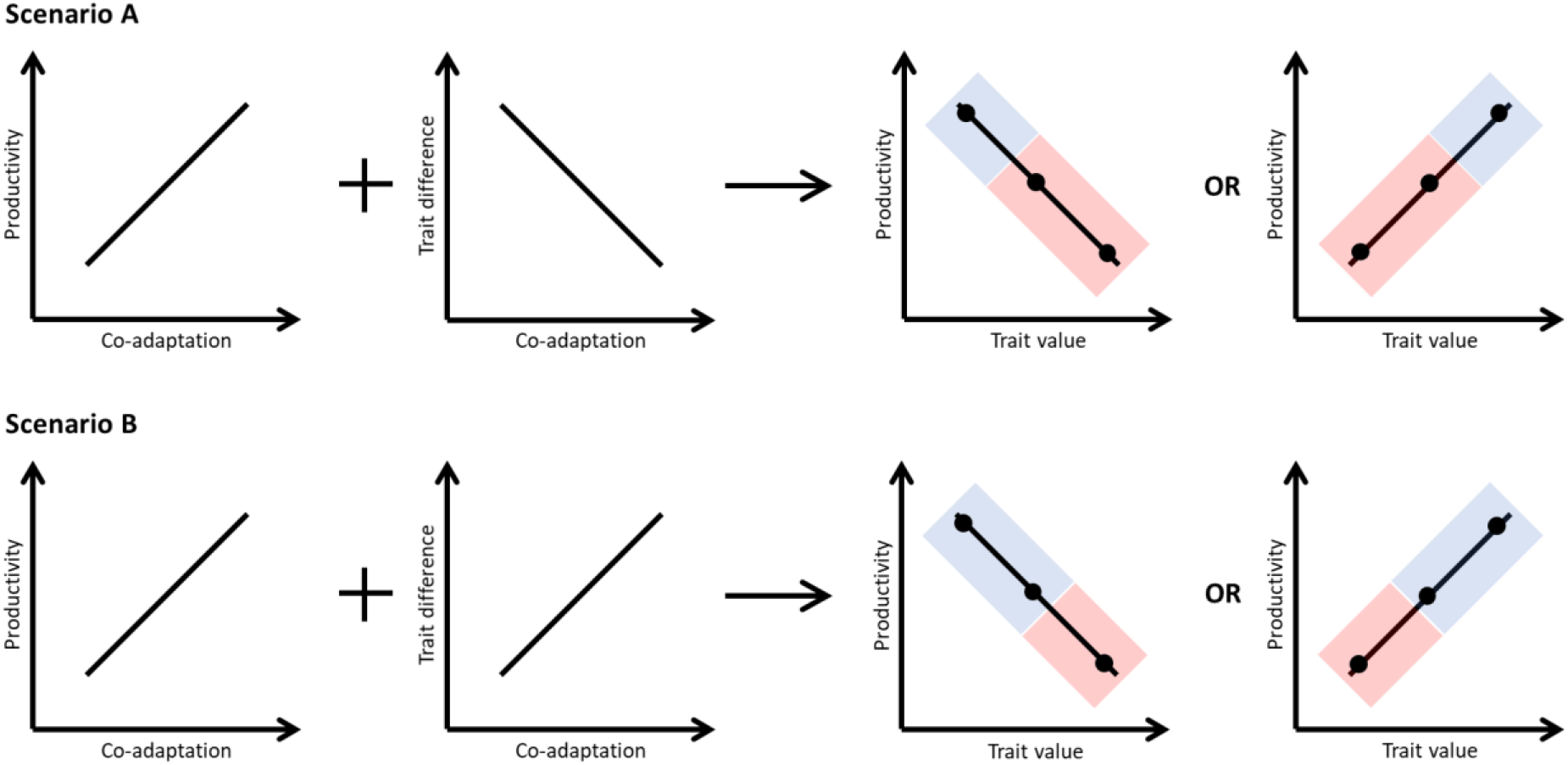
Schematic drawing of how co-adaption with enhanced productivity and trait convergence/divergence results in the selection of plants with high productivity (blue shaded area) and evolutive exclusion of plants with certain trait values (red shaded area). In scenario A, co-adaption with enhanced productivity and trait convergence results in the selection of plants with high productivity and extreme trait values and evolutive exclusion of plants with lower productivity and less extreme trait values. In scenario B, co-adaption with enhanced productivity and trait divergence results in the selection of plants with high and medium productivity, and extreme and medium trait values. The number of plants (black dots) with evolutive exclusion is larger in scenario A than scenario B.

### Conclusion

Our results suggest that *coadaptation* among plants might enhance ecosystem functioning (i.e. higher productivity). Nevertheless, there is still a lot of research needed to understand more about coevolution among plants. Especially interesting would be to identify on which functional traits coevolution is acting. These functional traits could again be targets for future breeding programs to sustainably enhance agricultural productivity.

## Acknowledgements

We thank Sandra González Sánchez, Francisco Ordiales Rosado, Jesús López-Angulo, Pilar Hurtado and Markus Bittlingmaier for their support in the field and Elisa Pizarro Carbonell from the Aprisco association for access to their research facilities. This work was funded by the ETH Zurich Research Grant ETH-28 19-2. The authors declare no competing interests.

## Authors contribution

AS planned and conducted the experiment, analysed the data, and wrote the manuscript. CS developed the idea, received the funding, planned the experiment and gave advise during the experiment, analysis and manuscript writing.

## Data availability statement

The data used in this study is available on Zenodo: https://doi.org/10.5281/zenodo.7620447.

