## Supplementary 1 for "Coadaptation of coexisting plants enhances productivity in an agricultural system"

**Table S1**. Overview of the planting scheme (2x2 Latin square). The shapes (circle and triangle) are different species, the shading are the different selection histories.

| **Culture** | **History composition** | |
| --- | --- | --- |
|  | **pure** | **mixed** |
| monoculture | 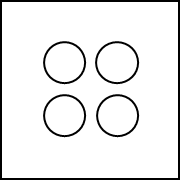 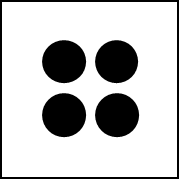 | 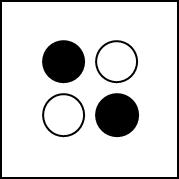 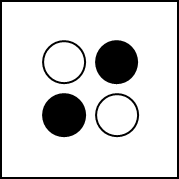 |
| mixture | 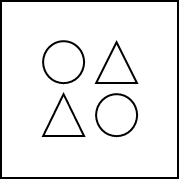 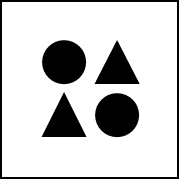 | 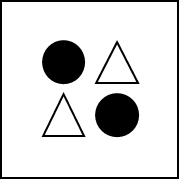 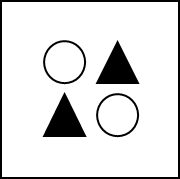 |

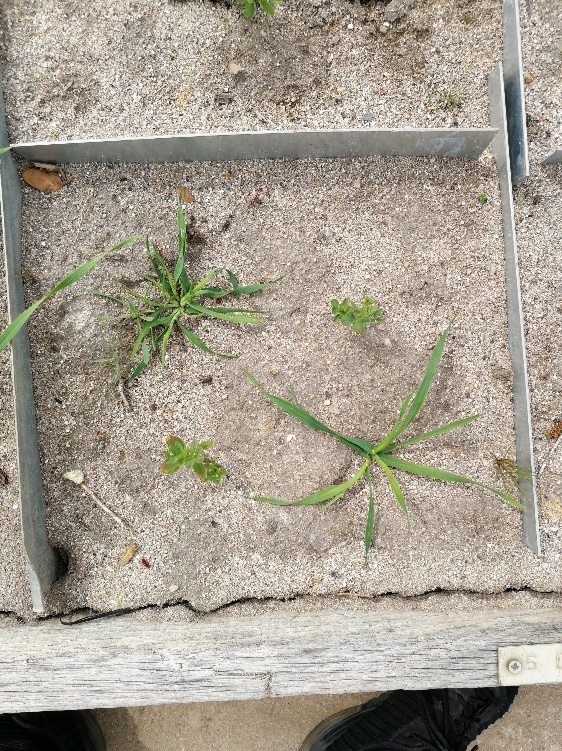

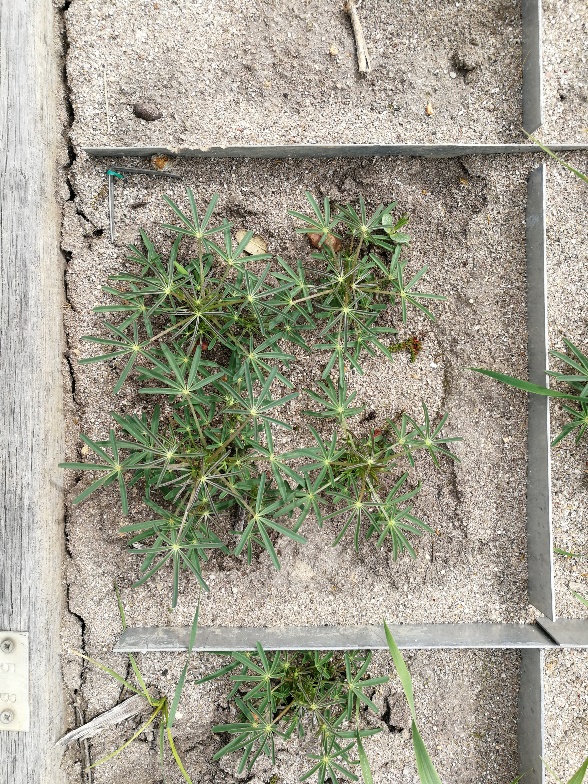

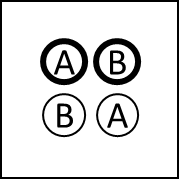

**b)**

**c)**

**a)**

**Fig. S1**. Examples of the planting scheme (2x2 Latin square). (**a**) Mixture of wheat (position a) and lentil (position b). (**b**) monoculture of lupin (position a and b). (**c**) The positions within the 2x2 Latin square. The individuals of the same species and selection history were always in the diagonal. The thick circles indicate individuals that were sampled during the functional trait measurement.

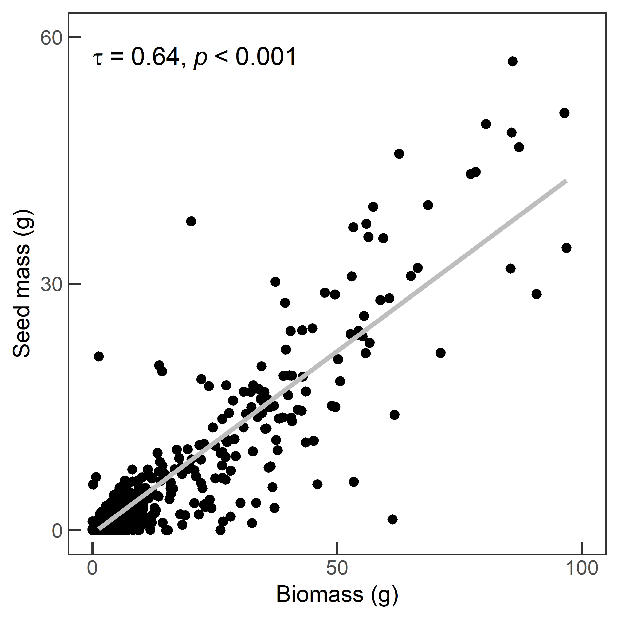

**Fig. S2**.Relationship between biomass and seed mass (individual level). The results from the Kendall correlation are also indicated.

a) community level

$$response \sim transplanted \left( a \right)+transplanted \left( b \right)+sowing \left( a \right)+sowing \left( b \right)+culture+familiarity+(1\left| spcomp \right)$$

used for: plot biomass, plot difference in functional traits

$$response \sim transplanted \left( a \right)+transplanted \left( b \right)+sowing \left( a \right)+sowing \left( b \right)+culture+familiarity+familiarity^2+(1\left| spcomp \right)$$

used for: plot seed mass

b) species level

$$\mathrm{response} \sim transplanted+sowing+culture+ species*selection history*history composition+\left( 1|plot \right)+\left( 1 \right|plot:position)+(1\left| spcomp \right)+(1|spcomp:hist)$$

used for: biomass, seed mass (zero-truncated), binominal seed mass, harvest index (zero-truncated)

$$\mathrm{response}\sim transplanted+sowing+culture+ species*selection history*history composition+\left( 1|plot \right)+(1\left| spcomp \right)+(1|spcomp:hist)$$

used for: coordinates of PC1 from functional traits

**Fig. S3**. Formulas of the linear mixed models and generalised linear mixed models.

**Table S2**.Type-III analysis of variance of the binominal seed mass (0=no seeds, 1=seeds) as response variable and the explanatory variables transplanted (yes vs no), sowing (first vs second), culture (monoculture vs mixture), species (oat, wheat, lentil, lupin, camelina or coriander), selection history (single vs community), history composition (pure vs mixed) and all possible interactions between species, selection history and history composition. The random terms were position within plot, species composition and the interaction species composition and selection history. DF: degrees of freedom; Chisq: Chi-square value; P: error probability. P-values in bold are significant at α = 0.05.

|  | DF | Chisq | P |
| --- | --- | --- | --- |
| transplanted | 1 | 33.77 | **<0.001** |
| sowing | 1 | 32.91 | **<0.001** |
| culture | 1 | 1.03 | 0.310 |
| species | 5 | 13.39 | **0.020** |
| selection history | 1 | 1.42 | 0.234 |
| history composition | 1 | 0.19 | 0.660 |
| species × selection history | 5 | 6.55 | 0.256 |
| species × history composition | 5 | 3.07 | 0.689 |
| selection history × history composition | 1 | 0.60 | 0.439 |
| species × selection history × history composition | 5 | 1.89 | 0.864 |

**Table S3**. Type I-analysis of variance of the response variables plot biomass (square-root transformed) and plot seed mass (log-transformed, zero-truncated) and the explanatory variables transplanted (position a and b), sowing (position a and b), culture (monoculture vs mixture), community history composition (pure single=1, mixed=2, pure community=3) and the community history composition second-degree polynomial (only in seed mass). The random term was the species composition. DF: degrees of freedom; DenDF: degrees of freedom of error term; F: probability distribution; P: error probability. P-values in bold are significant at α = 0.05.

|  | DF | DenDF | F | P |
| --- | --- | --- | --- | --- |
|  | plot biomass | | | |
| transplanted (position a) | 2 | 258.48 | 8.03 | **<0.001** |
| transplanted (position b) | 2 | 202.92 | 10.89 | **<0.001** |
| sowing (position a) | 2 | 247.92 | 21.80 | **<0.001** |
| sowing (position b) | 2 | 312.39 | 13.02 | **<0.001** |
| culture | 1 | 15.66 | 0.21 | 0.653 |
| community history composition | 1 | 306.84 | 5.28 | **0.022** |
|  | plot seed mass | | | |
| transplanted (position a) | 2 | 300.52 | 1.83 | 0.162 |
| transplanted (position b) | 2 | 289.22 | 9.22 | **<0.001** |
| sowing (position a) | 2 | 287.53 | 6.44 | **0.002** |
| sowing (position b) | 2 | 303.63 | 5.30 | **0.005** |
| culture | 1 | 15.21 | 0.55 | 0.471 |
| community history composition | 1 | 291.58 | 0.82 | 0.366 |
| community history composition  (second-degree polynomial) | 1 | 290.88 | 3.41 | 0.066 |

**Table S4**. Type I-analysis of variance of the response difference in plant height, specific leaf area (SLA) and leaf dry matter content (LDMC) and the explanatory variables culture (monoculture vs mixture), species (oat, wheat, lentil, lupin, camelina or coriander), selection history (single vs community), history composition (pure vs mixed) and all possible interactions between species, selection history and history composition. The random terms were species composition and the interaction species composition and selection history. DF: degrees of freedom; DenDF: degrees of freedom of error term; F: probability distribution; P: error probability. P-values in bold are significant at α = 0.05.

|  | DF | DenDF | F | P |
| --- | --- | --- | --- | --- |
|  | difference in plant height | | | |
| transplanted (position a) | 1 | 228.66 | 0.03 | 0.866 |
| transplanted (position b) | 1 | 222.41 | 0.03 | 0.865 |
| sowing (position a) | 1 | 76.98 | 1.20 | 0.277 |
| sowing (position b) | 1 | 165.53 | 2.42 | 0.122 |
| culture | 1 | 18.04 | 5.53 | **0.030** |
| community history composition | 1 | 228.08 | 0.00 | 0.977 |
|  | difference in SLA | | | |
| transplanted (position a) | 1 | 237.87 | 1.00 | 0.320 |
| transplanted (position b) | 1 | 235.97 | 0.40 | 0.528 |
| sowing (position a) | 1 | 94.35 | 0.06 | 0.803 |
| sowing (position b) | 1 | 174.31 | 0.73 | 0.965 |
| culture | 1 | 15.70 | 1.39 | 0.256 |
| community history composition | 1 | 226.14 | 8.37 | **0.004** |
|  | difference in LDMC | | | |
| transplanted (position a) | 1 | 242 | 1.95 | 0.163 |
| transplanted (position b) | 1 | 242 | 4.58 | 0.033 |
| sowing (position a) | 1 | 242 | 3.03 | 0.083 |
| sowing (position b) | 1 | 242 | 0.08 | 0.782 |
| culture | 1 | 242 | 0.01 | 0.906 |
| community history composition | 1 | 242 | 1.21 | 0.242 |

**Table S5**. Type I-analysis of variance of the response variables biomass (log-transformed), seed mass (log-transformed, zero-truncated) and the harvest index (square-root transformed, zero-truncated). The explanatory variables were transplanted (yes vs no), sowing (first vs second), culture (monoculture vs mixture), species (oat, wheat, lentil, lupin, camelina or coriander), selection history (single vs community), history composition (pure vs mixed) and all possible interactions between species, selection history and history composition. The random terms were position within plot, species composition and the interaction species composition and selection history. DF: degrees of freedom; DenDF: degrees of freedom of error term; F: probability distribution; P: error probability. P-values in bold are significant at α = 0.05.

|  | DF | DenDF | F | P |
| --- | --- | --- | --- | --- |
|  | biomass | | | |
| transplanted | 1 | 696.84 | 1335.13 | **<0.001** |
| sowing | 1 | 275.81 | 69.32 | **<0.001** |
| culture | 1 | 11.17 | 1.17 | 0.302 |
| species | 5 | 182.92 | 256.63 | **<0.001** |
| selection history | 1 | 12.03 | 1.33 | 0.271 |
| history composition | 1 | 294.66 | 0.65 | 0.420 |
| species × selection history | 5 | 39.29 | 0.58 | 0.713 |
| species × history composition | 5 | 595.82 | 3.72 | **0.003** |
| selection history × history composition | 1 | 612.02 | 1.57 | 0.210 |
| species × selection history × history composition | 5 | 609.49 | 2.30 | **0.043** |
|  | seed mass | | | |
| transplanted | 1 | 373.34 | 241.60 | **<0.001** |
| sowing | 1 | 152.89 | 69.72 | **<0.001** |
| culture | 1 | 10.61 | 1.67 | 0.224 |
| species | 5 | 48.00 | 124.83 | **<0.001** |
| selection history | 1 | 12.74 | 1.91 | 0.191 |
| history composition | 1 | 236.49 | 3.13 | 0.078 |
| species × selection history | 5 | 47.63 | 2.18 | 0.072 |
| species × history composition | 5 | 477.59 | 1.08 | 0.369 |
| selection history × history composition | 1 | 491.94 | 0.00 | 0.927 |
| species × selection history × history composition | 5 | 498.78 | 1.25 | 0.286 |
|  | harvest index | | | |
| transplanted | 1 | 601.70 | 86.43 | **<0.001** |
| sowing | 1 | 295.63 | 3.62 | 0.058 |
| culture | 1 | 24.57 | 3.04 | 0.094 |
| species | 5 | 115.43 | 44.14 | **<0.001** |
| selection history | 1 | 18.48 | 2.31 | 0.145 |
| history composition | 1 | 250.05 | 2.16 | 0.143 |
| species × selection history | 5 | 58.47 | 2.86 | **0.022** |
| species × history composition | 5 | 501.39 | 0.43 | 0.828 |
| selection history × history composition | 1 | 490.33 | 0.04 | 0.850 |
| species × selection history × history composition | 5 | 506.60 | 1.91 | 0.091 |

**Table S6**. Type I-analysis of variance of the response variable coordinates of PC1 from the functional traits and the explanatory variables transplanted (yes vs no), sowing (first vs second), culture (monoculture vs mixture), species (oat, wheat, lentil, lupin, camelina or coriander), selection history (single vs community), history composition (pure vs mixed) and all possible interactions between species, selection history and history composition. The random terms were species composition and the interaction species composition and selection history. DF: degrees of freedom; DenDF: degrees of freedom of error term; F: probability distribution; P: error probability. P-values in bold are significant at α = 0.05.

|  | DF | DenDF | F | P |
| --- | --- | --- | --- | --- |
| transplanted | 1 | 456.06 |  | **<0.001** |
| sowing | 1 | 457.25 |  | **<0.001** |
| culture | 1 | 234.70 |  | 0.187 |
| species | 5 | 469.92 |  | **<0.001** |
| selection history | 1 | 429.25 |  | 0.561 |
| history composition | 1 | 234.03 |  | 0.500 |
| species × selection history | 5 | 469.20 |  | 0.635 |
| species × history composition | 5 | 468.93 |  | 0.666 |
| selection history × history composition | 1 | 466.34 |  | **0.013** |
| species × selection history × history composition | 5 | 465.71 |  | 0.743 |

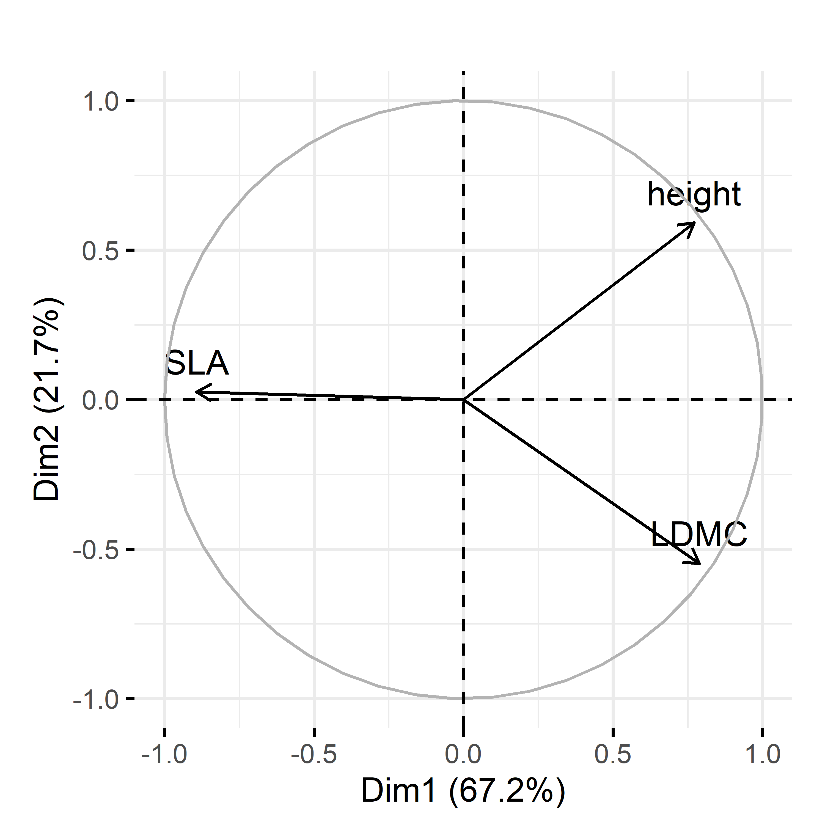

**Fig. S4**. The first two dimensions of the principal component analysis with the three functional traits plant height (height), specific leaf area (SLA) and leaf dry matter content (LDMC).

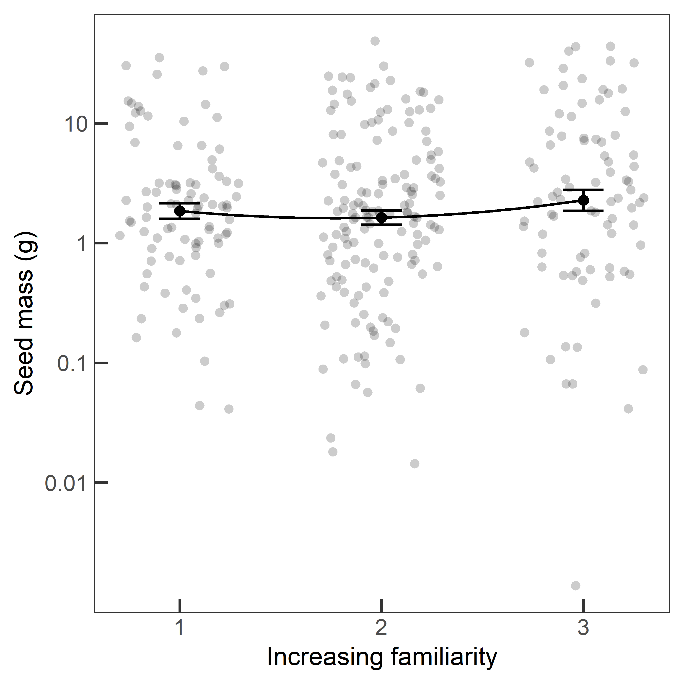

**Fig. S5**. Plot seed mass (log-scale, zero-truncated) of communities with increasing familiarity (pure *single*=1, *mixed*=2, pure *community*=3). Shown are the single data points, the mean ± standard error of the mean and the regression line (formula: y~x+x^2^). For statistics see Table S3.

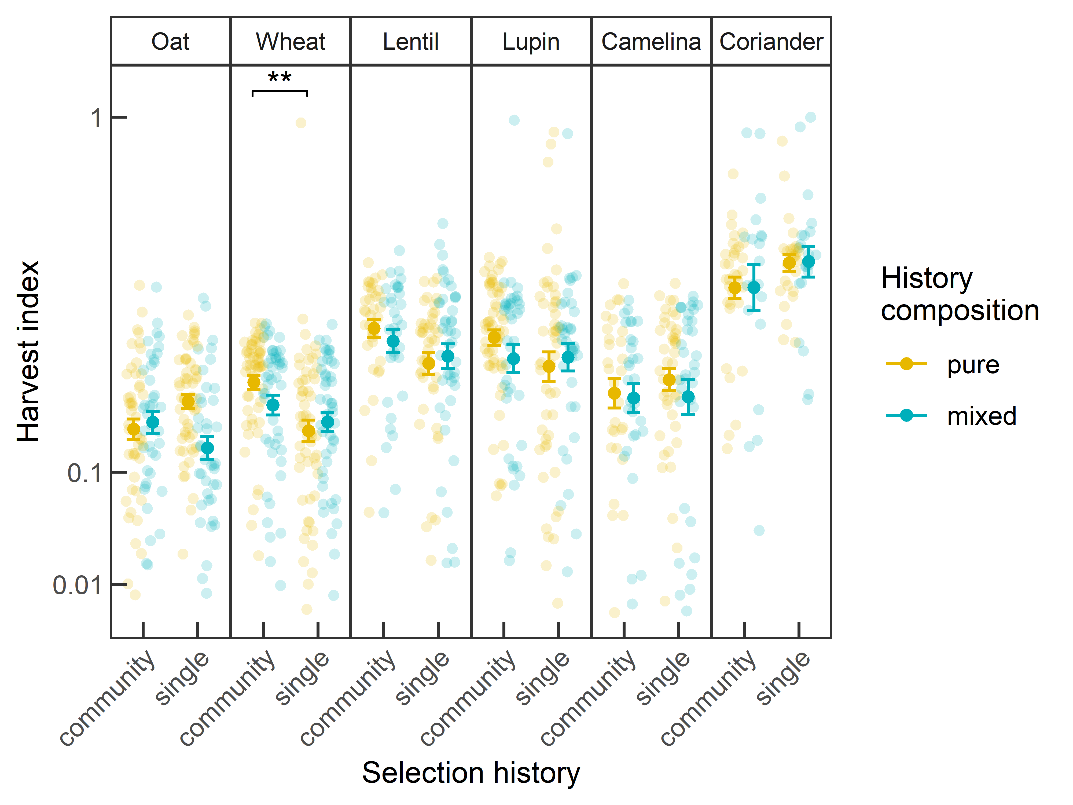

**Fig. S6**. Harvest index (square-root scale, zero-truncated) of plants from either *community* or *single* selection history and planted in either *pure* (yellow) or *mixed* (blue) history composition. Shown are the single data points and the mean ± standard error of the mean (point and error bar). Significant differences to test the hypotheses (Fig. 1b) are indicated with brackets and asterisk (post-hoc, estimated marginal means). Asterisks correspond to *: P<0.05; **: P<0.01; ***: P<0.001. For statistics see Table S5.

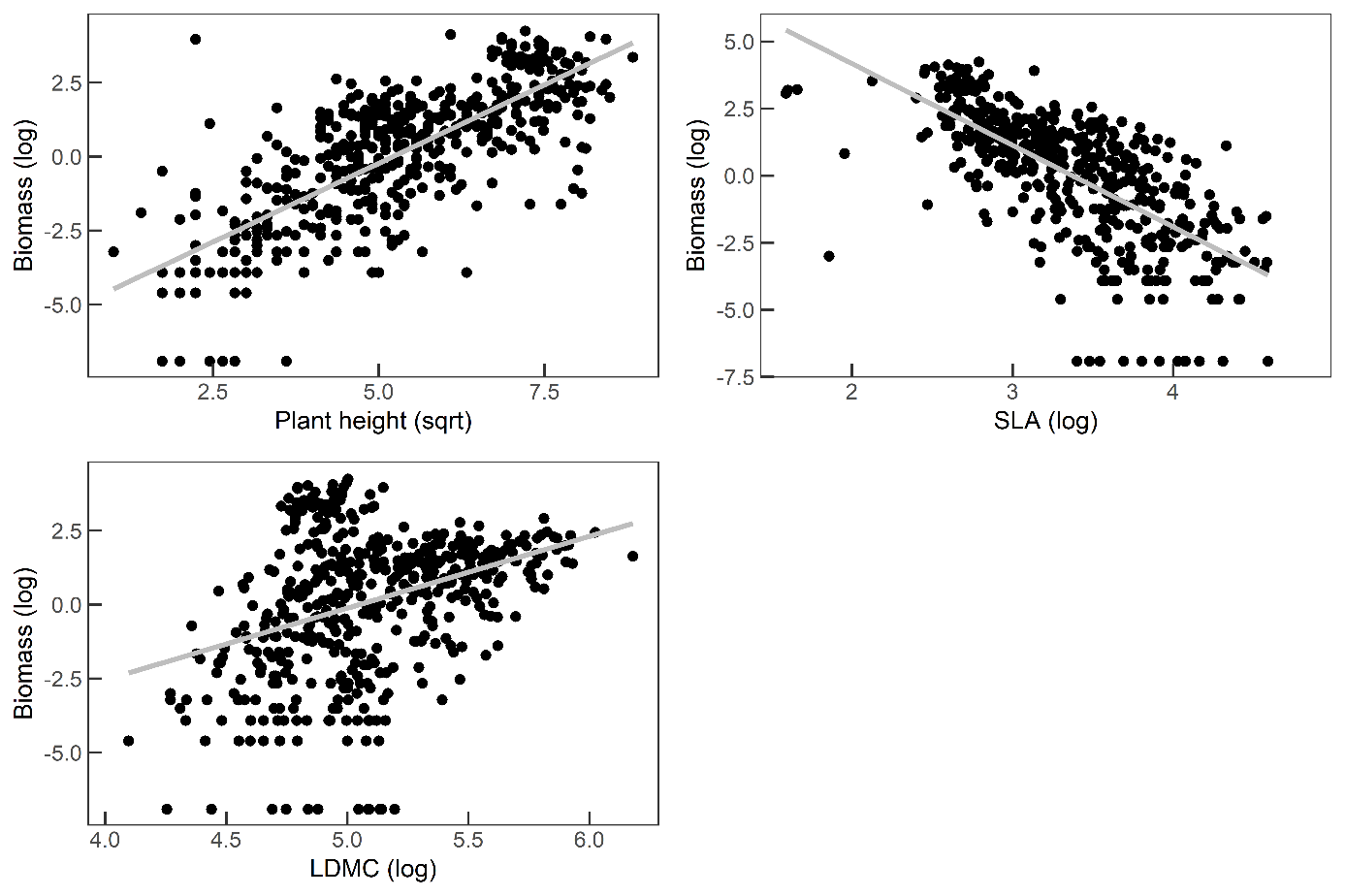

**Fig. S7**. Relationship between biomass and the functional traits plant height, specific leaf area (SLA) and leaf dry matter content (LDMC). Variables are either log- (biomass, SLA and LDCM) or square-root transformed (plant height).
